# Mangrove propagules are limited in their capacity to disperse across long distances

**DOI:** 10.1101/2023.01.30.526169

**Authors:** Natasha R. Granville, Cristina Banks-Leite

## Abstract

Mangroves are subject to rapid and large-scale habitat changes which threaten their unique genetic diversity and provision of critically important ecosystem services. Habitat fragmentation reduces connectivity which can impair dispersal and lead to genetic isolation. However, it is unclear whether fragmentation could impact mangrove genetic isolation, as mangrove propagules can disperse long distances. Here, we conducted a meta-analysis of studies reporting a correlation between geographic distance and genetic distance in mangrove plants. From the 22 studies that met the inclusion criteria, we found a significant isolation-by-distance effect; geographic distance was significantly associated with Nei’s genetic distance and F_ST_. Our results show that mangrove propagules may be limited in their capacity to disperse across long distances, which highlights the importance of maintaining close proximity between habitat patches and reducing habitat fragmentation.

## Introduction

Habitat loss is known to impact dispersal across habitats, which is vital for maintaining genetic diversity and for range shifts in response to environmental changes (Van der Stocken et al. 2019a). Understanding the relationship between spatial and genetic structuring of populations is fundamental for developing effective conservation strategies that maintain habitat connectivity (Durrant et al. 2014, Taylor et al. 2021, Wright et al. 2015), and this understanding is contingent on characterising dispersal capabilities.

Mangrove forests are intertidal wetlands, found along coastlines in tropical, sub-tropical and warm-temperate climates (Bryan-Brown et al. 2020). Globally, there are two main hotspots of mangrove biodiversity; the Indo-West Pacific contains approximately 54 mangrove species, and the Atlantic-East Pacific contains approximately 17 mangrove species (Tomlinson 2016). Between 2000 and 2012, mangrove forests were lost at an average rate of 0.18 % per year in Southeast Asia (Richards & Friess 2016). Mangroves are threatened at the local-and regional-scales by aquaculture, agriculture, urban development and pollution, while simultaneously facing the broader scale threats of sea level rise and climate change. Mangrove forests are of exceptionally high ecological and economic value (estimated $194,000 per hectare per year, Costanza et al. 2014) as they provide unique genetic diversity, and crucial ecosystem services (Bryan-Brown et al. 2020; Mantiquilla et al. 2021). The provision of these services depends on the size and arrangement of forest patterns (Bryan-Brown et al. 2020) Even in areas with low rates of mangrove loss, there is a global trend towards ubiquitous fragmentation, which can pose a threat to mangrove biodiversity (Bryan-Brown et al. 2020). Therefore, there is a need to quantify the impact of fragmentation on mangrove biodiversity.

While it is generally accepted that habitat loss has negative effects on biodiversity, the effects of habitat fragmentation *per se* (independent of habitat loss) are more variable and context-dependent (Andrén 1994, Fahrig 2003). On the one hand, Wilcove et al’s 1986 definition of habitat fragmentation as a process whereby *“a large expanse of habitat is transformed into a number of smaller patches of smaller total area, isolated from each other by a matrix of habitats unlike the original”* (Wilcove 1986, as cited in Fahrig 2003) implies that fragmentation leads to increased isolation of patches (Fahrig, 2003). On the other hand, fragmentation *per se* could theoretically result in many small patches that are less isolated from each other, assuming that dispersal across the matrix is possible (Fahrig 2003, Healey & Hovel 2004). In mangroves, however, the matrix between habitat patches is composed of water, which can be easily traversed by propagules and human-modified land use, which cannot be colonised by mangroves.

It is therefore important to understand the impacts of geographic distance on mangrove propagule dispersal. Buoyant mangrove propagules have the capacity for long-distance dispersal by water because they remain viable for extended periods of time, and can drift in ocean currents (Binks et al. 2019). This could be expected to lead to high connectivity between habitat patches, which could mean that increasing distance between patches would have little effect on genetic isolation. However, field studies have shown that mangrove propagules tend not to disperse far from their release point, leading to patterns of isolation-by-distance (Binks et al. 2019, Clarke 1993, Yan et al. 2016,Van der Stocken et al. 2015). Even in species with highly dispersive life history features, abiotic factors like ocean currents, coastal topography and habitat discontinuities can affect the realised dispersal distance (Cowen & Sponaugle 2009, Selkoe et al. 2016, Binks et al. 2019). Propagules of *Avicennia marina*, the world’s most widely distributed mangrove species, can remain viable in water for several months (Rabinowitz 1978). However, their obligate dispersal period, during which they float before developing roots for anchoring, is only around one week (Rabinowitz 1978). Once they land on suitable substrate, they can root and grow rapidly because many taxa exhibit viviparity, where the seed germinates while attached to the parent plant (Clarke 1993, Ng & Sivasothi 2001). These traits, combined with the risks of dispersal, could explain why mangrove propagules might tend to favour establishment near their release point (Clarke 1993, Binks et al. 2019).

We conducted a meta-analysis to quantify the relationship between geographic distance and genetic distance in mangrove plant communities to better understand how far mangrove propagules disperse, with the aim of providing insight into the effect of patch isolation on genetic isolation in mangroves. Measures of genetic distance such as Nei’s genetic distance (Nei 1972) and F_ST_ (Wright 1950) provide useful insights into the genetic structure of a habitat (Bohonak 1999). If mangrove patches are more isolated from each other and propagules cannot disperse between them, there will likely be a larger pairwise genetic distance between these patches (Jaquiéry et al. 2011).

## Methods

### Systematic literature search and inclusion criteria

In May 2021, we conducted an extensive search of the relevant literature, following the PRISMA (Preferred reporting items for systematic review and meta-analysis) statement which provides a standardised framework for reporting systematic reviews and meta-analyses (Moher et al. 2009). The following search terms were used in Web of Science and Scopus:

1. mangrove AND fragment* AND (‘genetic diversity’ OR ‘genetic differentiation’)
2. mangrove AND connect* AND (‘genetic diversity’ OR ‘genetic differentiation’)
3. mangrove AND isolat* AND (‘genetic diversity’ OR ‘genetic differentiation’)

An initial search, after removing duplicates, yielded 199 papers, which we screened for eligibility. This resulted in the exclusion of 148 non-relevant papers and 4 papers that we could not access. From the 47 papers that remained, we selected papers that reported the results of a Mantel test for the correlation between matrices of untransformed Euclidean geographic distance on Nei’s genetic distance (11 papers, representing 13 case studies) or untransformed Wright’s F_ST_ (8 papers, representing 9 case studies). Nei’s genetic distance and F_ST_ were chosen as measures of genetic diversity because they were the most commonly reported so this helped maximise the sample size. We expect that our sample size of 22 studies is large enough to provide sufficient power to make meaningful conclusions (Jackson & Turner 2017). The analysis of the effect of geographic distance on Nei’s genetic diversity is henceforth referred to as ‘Nei’s meta-analysis’ and the analysis of the effect of geographic distance on Wright’s F_ST_ is henceforth referred to as ‘F_ST_ meta-analysis’

### Statistical analysis

For the effect size, we extracted Pearson’s *r* value from the reported Mantel test in each paper. Where R^2^ was reported, it was converted to *r* by taking the square root (this was done in 2 papers for the Nei’s meta-analysis and 8 papers for the F_ST_ meta-analysis). Each study was weighted according to the following formula developed by Reed & Frankham (2003) specifically for meta-analyses of genetic diversity: 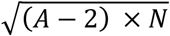, where *A* = number of populations and *N* = number of individuals for each paper.

Analyses were conducted using the metafor (Viechtbauer 2010) and robumeta (Fisher et al. 2017) packages in R version 4.2.1 (R Core Team 2022). We used a random-effects model because studies were drawn from different populations. For the analysis, Pearson’s *r* was transformed to Fisher’s *Z* to ensure normal distribution. After performing meta-analytic calculations, Fisher’s *Z* was converted back to Pearson’s *r* for reporting summary effect sizes (Quintana 2015).

The Q-statistic was calculated to assess heterogeneity among studies. The Q-statistic is the ratio of observed variation to within-study variance. It evaluates the null hypothesis that all studies are examining the same effect (Quintana 2015). Different studies used different mangrove plant species and different molecular markers to assess genetic variation (Table S1). To assess the effect of this, we fitted separate meta-analytic models that moderated for the effects of species and marker, respectively. To account for effect size dependency resulting from the same study reporting multiple effect sizes (2 papers in the Nei’s meta-analysis and 1 paper in the F_ST_ meta-analysis), we used robust variance estimation as this is appropriate for meta-analyses with less than 40 studies and does not assume knowledge of within-study correlations (Quintana 2015). We used Egger’s regression test to assess publication bias by testing for funnel plot asymmetry. Publication bias is the phenomenon whereby studies with larger effect sizes are more likely to be published, and therefore included in the meta-analysis (Quintana 2015).

## Results

### Included studies

There were 13 case studies for Nei’s genetic distance and 9 case studies for F_ST_ (Table S1, Figure 1). Egger’s regression test for funnel plot asymmetry showed no effect of publication bias (Nei’s meta-analysis: *z* = 0.158, p= 0.875. F_ST_ meta-analysis: *z* = 1.31, p = 0.192. Figure S1).

**Figure 1.**
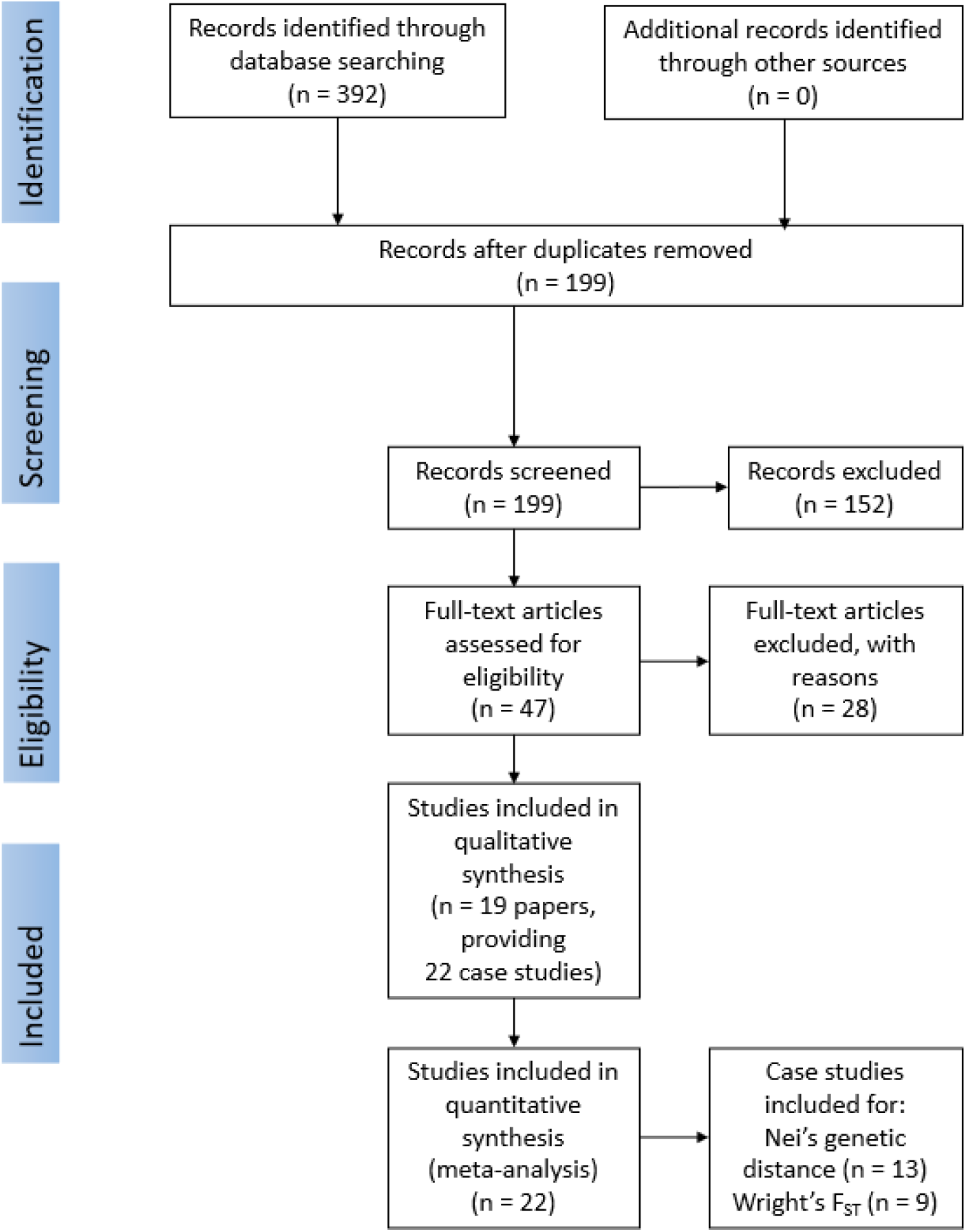
PRISMA flowchart (Moher et al. 2009) showing the sequence of selection of papers for meta-analysis of the effects of geographic distance on genetic distance.

### Effect sizes

We found a significant association between geographic distance and Nei’s genetic distance (estimated model coefficient = 0.37, 95 % CI = 0.14 -0.56. *z* = 3.07, p = 0.002. Figure 2A), which was not changed when accounting for effect size dependency by robust variance estimation (estimated model coefficient = 0.39, 95 % CI = 0.12 – 0.66). We also found a significant association between geographic distance and F_ST_ (estimated model coefficient = 0.63, 95 % CI = 0.41 – 0.78. *z* = 4.77, p < 0.0001. Figure 2b). This model coefficient was not changed significantly when accounting for effect size dependency by robust variance estimation (estimated model coefficient = 0.75, 95 % CI = 0.38 – 1.11).

**Figure 2.**
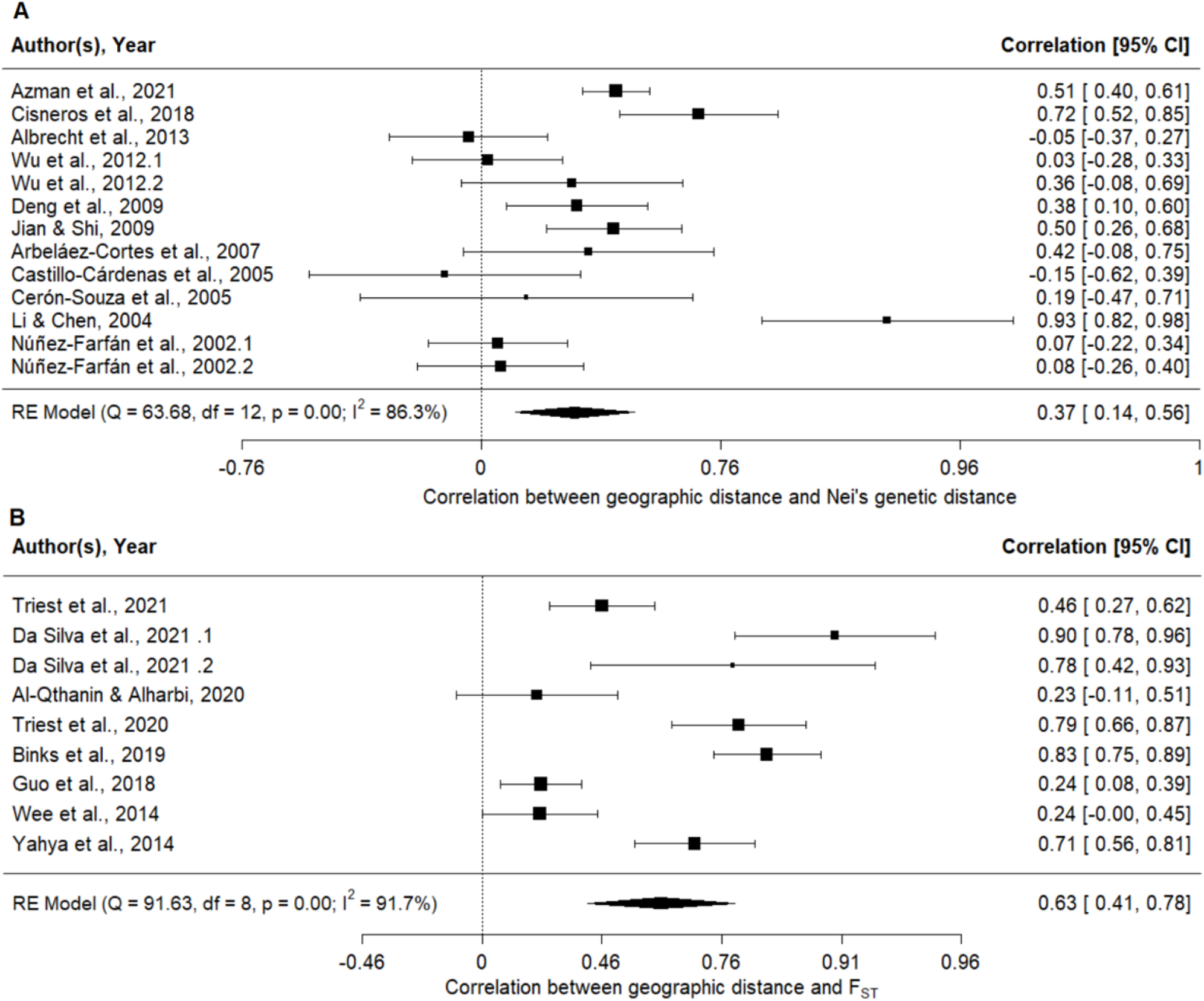
Effect of geographic distance on (A) Nei’s genetic distance and (B) genetic distance measured by F_ST_ for the original unadjusted models, with Fisher’s *Z* transformed back to Pearson’s *r*. The polygon at the bottom of each plot shows the estimated model coefficient and its bounds represent the 95 % confidence intervals (CIs). Each squared point corresponds to a different study, as labelled. The size of the square corresponds to the contribution of the study to the estimated model coefficient.

### Heterogeneity and moderator analysis

There was significant heterogeneity among the included studies (Nei’s meta-analysis: Q = 63.7, df = 12, p < 0.0001. F_ST_ meta-analysis Q = 91.6, df = 8, p < 0.0001). Li & Chen (2004) contributed disproportionately to the overall heterogeneity in Nei’s meta-analysis. Therefore, for the purpose of comparison, a separate random-effects model was fitted to the same data set but excluding Li & Chen (2004). This reduced the summary effect size from 0.37 (95 % CI = 0.14 -0.56) to 0.29 (95 % CI = 0.12 -0.45). The overall heterogeneity was reduced, but there was still significant heterogeneity (Q = 40.4, df = 11, p <0.0001). Since excluding this study did not significantly remove the heterogeneity, all other analyses include this study.

To determine the source of heterogeneity, we conducted mixed-effects moderator analyses with taxon and marker as separate moderator variables. The type of genetic marker significantly moderated the correlation between geographic distance and F_ST_ (QM = 12.0, df = 3, p = 0.0075), but not Nei’s genetic distance (QM = 3.63, df = 4, p = 0.46). Whereas differences in the taxon investigated did not significantly moderate either of these correlations (Nei’s genetic distance: QM = 12.5, df = 7, p = 0.086. F_ST_: QM = 7.50, df = 6, p = 0.28).

## Discussion

Our global meta-analysis showed a significant correlation between geographic distance and genetic distance in mangrove plant communities. This isolation-by-distance effect (Wright 1943) could suggest that mangrove plants are limited in their capacity to disperse across habitat patches. This is consistent with the conclusions made by Binks et al. (2019) that habitat discontinuities lead to reduced gene flow between patches because mangrove propagules tend not to disperse far from their release point, likely due to a combination of abiotic factors, viviparity and short obligate dispersal period (Clarke 1993, Binks et al. 2019). Maintaining gene flow, which is critical for long-term population persistence (Salm et al. 2000, Wright et al. 2015), will depend on maintaining proximity among habitat patches, especially under conditions of habitat transformation which threaten mangrove biodiversity.

Isolation-by-distance indicates that spatial structure and genetic structure are highly correlated and suggests that dispersal limitation may be important in mangrove communities. Dispersal is essential for enabling sessile organisms, such as plants, to move away from unfavourable conditions if they are unable to adapt to such conditions (Kinlan & Gaines 2003). Isolation-by-distance suggests that these important adaptive responses are constrained by natural dispersal mechanisms (Sexton et al. 2014). Though not explicitly considered in this study, isolation-by-environment and isolation-by-resistance are relevant to understanding the factors underpinning the genetic structure of mangrove populations. Isolation-by-resistance includes environmental factors, such as land-use changes and biogeographic barriers, that affect the ability of propagules to disperse between patches of suitable habitat. This can affect the genetic structure of populations by modulating the impact of geographic distance on genetic distance (Wang & Bradburd 2014). Furthermore, environmental factors such as habitat heterogeneity can affect the likelihood of gene flow among populations; and these isolation-by-environment factors can act in combination with geographic distance to drive the genetic structure of populations (McRae 2006).

If mangrove propagules are limited in their dispersal capabilities, populations in habitat patches are more likely to become isolated from each other. This could result in a meta-population structure with smaller populations that are more vulnerable to demographic and environmental stochasticity (Lande 1993). Furthermore, the apparent dispersal limitation of mangrove propagules could limit the potential for mangroves to shift their distributional ranges to track changing climatic conditions (Van der Stocken et al. 2022). While we recognise that dispersal depends on several biotic and abiotic factors that affect the release, transport and establishment of propagules (Van der Stocken et al. 2019b), the isolation-by-distance effect shown here highlights the importance of geographic distance in constraining gene flow and suggests that mangrove propagules are limited in their tendency to disperse across long distances. This may be exacerbated by future climatic warming and rising sea levels. Recent analysis by Van der Stocken et al. (2022) indicates that mangrove propagules in fresher and warmer oceans are likely to have increased rates of sinking, which reduces the likelihood of long distance dispersal, especially for widespread mangrove species with dense propagules such as *Avicennia marina* (Van der Stocken et al. 2022). Therefore, when conserving and managing mangroves, the importance of maintaining close proximity between habitat fragments should be considered.

Our results indicate that the genetic structure of mangrove communities is dependent on spatial structure. Existing efforts to restore deforested mangrove forests often involve artificial movement of propagules (Vanderklift et al. 2020) which might be relevant for assisting dispersal if this is needed to maintain connectivity. Furthermore, for protected area networks to successfully maintain landscape connectivity, the size and arrangement of these networks should reflect the dispersal capabilities of the inhabiting species (Durrant et al. 2014, Shanks et al. 2003). Therefore, optimal design of protected area networks requires knowledge of effective dispersal distances. While the present study does not address exact distances, our results suggest that the realised dispersal capabilities of mangrove propagules depend heavily on the geographic distances across which they are dispersing. This emphasises the need for future studies to quantify effective dispersal distances in mangroves and consider how mangrove dispersal could be affected by habitat change.

## Acknowledgements

We thank two anonymous reviewers for helpful comments on an earlier version of the manuscript.

