## Supplementary material for "Mangrove propagules are limited in their capacity to disperse across long distances": Figure S1; Figure S2; Table S1.

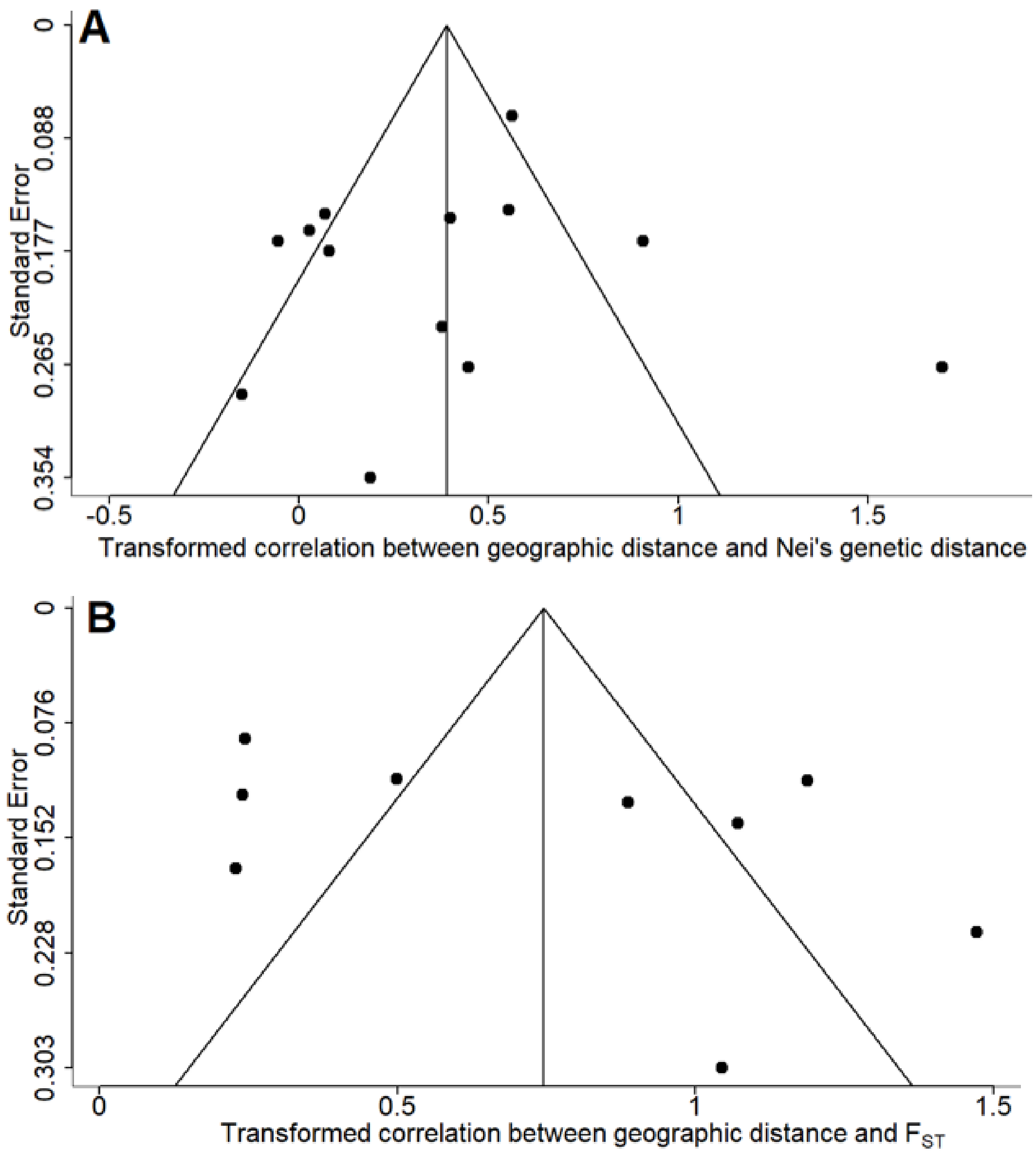


Figure S1. Funnel plots for Fisher's *z* transformed correlation coefficient (*x* axis) against study weight (standard error on a reverse scale, *y* axis) for (A) Nei's meta-analysis and (B) $F_{ST}$ meta-analysis.

Table S1. Studies included in the meta-analysis of the effect of geographic distance on (A) Nei's genetic distance (B) genetic distance measured by $F_{ST}$. The effect size is Pearson's *r* value, and the sample weight is according to the formula developed by Reed and Frankham 2003 (see methods). Taxon and genetic marker are moderator variables. For the genetic markers AFLP refers to Amplified Fragment Length Polymorphism and ISSR refers to Inter Simple Sequence Repeats. The full citations for the studies including the DOIs can be found in the reference list.

| **Study ID** | **Citation** | **Effect size** | **Number of individuals** | **Number of populations** | **Sample weight** | **Taxon** | **Genetic marker** | **Region** |
|---|---|---|---|---|---|---|---|---|
| **(A) Nei's genetic distance:** | | | | | | | | |
| **1** | Azman et al. 2020 | 0.51 | 1,120 | 39 | 204 | *Rhizophora apiculata* | Microsatellites | Malaysia |
| **2** | Cisneros-de la Cruz et al. 2018 | 0.72 | 130 | 13 | 38 | *Rhizophora mangle* | Microsatellites | Yucatan peninsula |
| **3** | Albrecht et al. 2013 | -0.05 | 130 | 13 | 38 | *Rhizophora mangle* | AFLP | Florida and Caribbean |
| **4** | Wu et al. 2012 | 0.03 | 220 | 10 | 42 | *Derris trifoliata* | ISSR | Southern China (mainland) |
| **5** | Wu et al. 2012 | 0.36 | 142 | 5 | 21 | *Derris trifoliata* | ISSR | Southern China (Hainan Island) |
| **6** | Deng et al. 2009 | 0.38 | 202 | 13 | 47 | *Aegiceras corniculatum* | AFLP | South China, Malay Peninsula, Sri |

| | | | | | | | | |
|---|---|---|---|---|---|---|---|---|
| | | | | | | | | Lanka, and North Australia |
| **7** | Jian & Shi 2009 | 0.50 | 234 | 13 | 51 | *Heritiera littoralis* | AFLP | China, Japan and Thailand |
| **8** | Arbeláez-Cortes et al. 2007 | 0.42 | 92 | 5 | 17 | *Rhizophora mangle* | Microsatellites | Colombian Pacific |
| **9** | Castillo-Cárdenas et al. 2005 | -0.15 | 57 | 6 | 15 | *Pelliciera rhizopborae* | AFLP | Colombian Pacific |
| **10** | Cerón-Souza et al. 2005 | 0.19 | 56 | 4 | 11 | *Avicennia germinans* | AFLP | Colombian Pacific |
| **11** | Li & Chen 2004 | 0.93 | 100 | 5 | 17 | *Sonneratia alba* | ISSR | Hainan Island, China |
| **12** | Núñez-Farfán et al. 2002 | 0.07 | 400 | 8 | 49 | *Rhizophora mangle* | Allozymes | Atlantic coast of Mexico |
| **13** | Núñez-Farfán et al. 2002 | 0.08 | 300 | 6 | 35 | *Rhizophora mangle* | Allozymes | Pacific coast of Mexico |
| | **(B) Wright's $F_{ST}$:** | | | | | | | |
| **1** | Triest et al. 2021 | 0.46 | 670 | 12 | 82 | *Avicennia marina* | Microsatellites | Kenya and Tanzania |

| | | | | | | | | |
|---|---|---|---|---|---|---|---|---|
| **2** | Da Silva et al. 2021 | 0.90 | 77 | 10 | 25 | *Avicennia saurian* | SNP | South American coast |
| **3** | Da Silva et al. 2021 | 0.78 | 48 | 6 | 14 | *Avicennia germinans* | SNP | South American coast |
| **4** | Al-Qthanin & Alharbi 2020 | 0.23 | 193 | 9 | 37 | *Avicennia marina* | Microsatellites | Farasan archipelago, the Red Sea, Saudi Arabia |
| **5** | Triest et al. 2020 | 0.79 | 457 | 8 | 52 | *Avicennia marina* | Microsatellites | Gazi Bay, Kenya |
| **6** | Binks et al. 2019 | 0.83 | 336 | 21 | 80 | *Avicennia marina* | SNP | Western Australia |
| **7** | Guo et al. 2018 | 0.24 | 418 | 47 | 137 | *Excoecaria agallocha* | Chloroplast DNA | Indo-West Pacific |
| **8** | Wee et al. 2014 | 0.24 | 432 | 13 | 69 | *Rhizophora mucronata* | Microsatellites | Southeast Asia |
| **9** | Yahya et al. 2014 | 0.71 | 312 | 15 | 64 | *Rhizophora apiculata* | Microsatellites | Greater Sunda Islands of Indonesia |

Figure S2: Map of locations of study sites included in the meta-analysis.

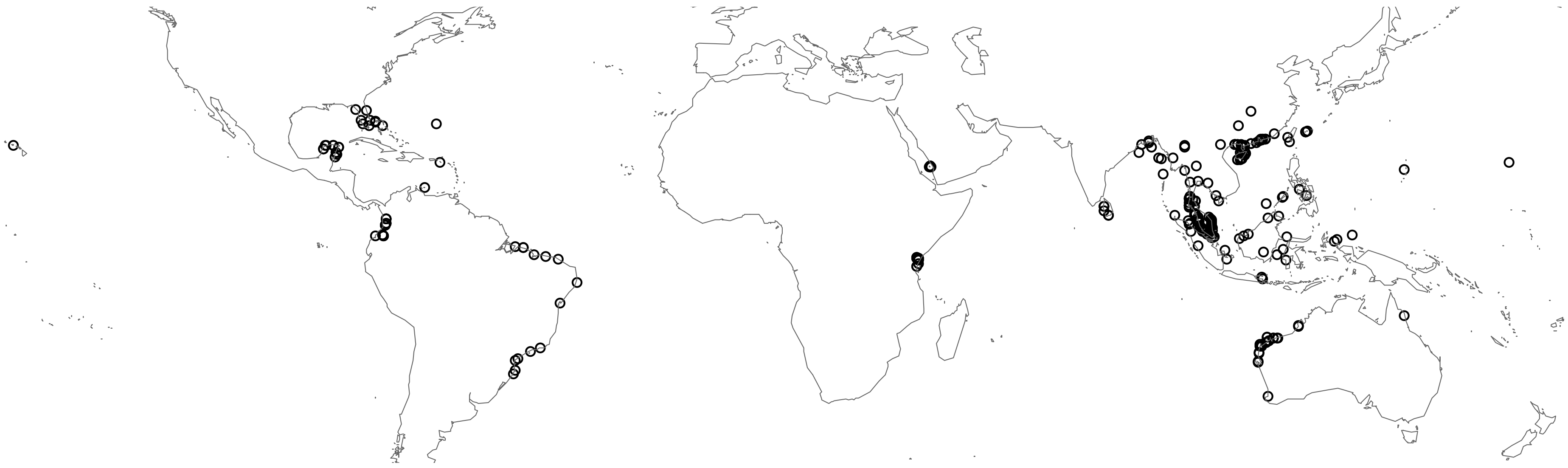
